# Loss of F-box Motif Encoding Gene SAF1 and RRM3 Together Leads to Synthetic Growth Defect and Sensitivity to HU, MMS in *S.cerevisiae*

**DOI:** 10.1101/636902

**Authors:** Meenu Sharma, V. Verma, Narendra K Bairwa

**Author notes:** To whom correspondence may be addressed /285634.

## Abstract

Unearthing of novel genetic interaction which leads to synthetic growth defects due to inactivation of genes are needed for applications in precision medicine. The genetic interactions among the molecular players involving different biological pathways need to be investigated. The SAF1 gene of *S.cerevisiae* encodes for a protein product which contain N-terminal F-box motif and C-terminal RCC1 domain. The F-box motif interacts with Skp1subunit of the SCF-E3 ligase and C-terminus with Aah1 (adenine deaminase) for ubiquitination and subsequent degradation by 26S proteasome during phase transition from proliferation state to quiescence phase due to nutrient limitation stress. The replication fork associated protein Rrm3 of *S.cerevisiae* belongs to Pif1 family helicase and function in removal of the non-histone proteins during replication fork movement. Here we have investigated the genetic interaction among both the genes (SAF1 and RRM3) and their role in growth fitness and genome stability. The single and double gene knockout strains of SAF1and RRM3 genes was constructed in BY4741 genetic background and checked for the growth fitness in presence of genotoxic stress causing agents such as hydroxyurea and methyl methanesulfonate. The strains were also evaluated for nuclear migration defect by DAPI staining and for HIS3AI marked Ty1 retro-transposition. The *saf1Δrrm3Δ* showed the extremely slow growth phenotype in rich medium and sensitivity to genotoxic agents such as HU and MMS in comparison to single gene mutant (*saf1Δ*, *rrm3Δ*) and WT cells. The *saf1Δrrm3Δ* also showed the defects in nuclear migration as evident by multi-nuclei phenotype. The *saf1Δrrm3Δ* also showed the elevated frequency of Ty1 retro-transposition in JC2326 background in comparison to either *saf1Δ or rrm3Δ*. Based on these observations we report that thatSAF1 and RRM3 functions in parallel pathway for growth fitness and stability of the genome.

## Introduction

The proliferation of *Saccharomyces cerevisiae* cells from in and out of quiescence phase due to nutrients stress or availability remains an active research area for both basic research and biotechnological purpose. The phase transition process is dependent on the ubiquitin proteasome system as it was reported that ubiquitin is required for survival during starvation (1).The SCF E3-ligase component Saf1, of *S.cerevisiae* which is an F-box motif containing protein recruits the Aah1 for proteasoaml mediated degradation upon nutrient deprivation (2). *The AAH1* gene of *S.cerevisiae*, encodes adenine deaminase (Aah1) which converts adenine to hypoxanthine. The nutrient deprivation condition induces stress which leads to cell enter into the quiescence phase. Besides the Aah1, the serine protease B (PRB1), protease C (PRC1), and Ybr139w, vacuolar origin proteins also have been reported as Saf1 targets for proteasome mediated degradation (3, 4).The serine proteases of the vacuole plays a major role during starvation of a cell.

The Pif1 family helicases are well conserved from yeast to humans. In *S. cerevisiae*, RRM3 gene encoded product Rrm3 belongs to Pif1 family, shows the 5-3’ helicase activity during replication fork movement (5, 6). Rrm3 is involved in the chromosome replication and travels along the replication fork (7).The activity of the Rrm3 counteracted by the replication fork protection complex, Tof1/Csm3 (8). In the absence of Rrm3, replication fork pausing occurs at the ITS1 region of r-DNA, telomeric regions, and difficult to replicate sites in the chromosome (8, 9). Rrm3 was first identified as suppressor of recombination in the repeat region(10).The loss of RRM3 results in the r-DNA circle formation which suggests its role in maintenance of the r-DNA stability. The rrm3 mutant showed the synthetic fitness defect when combined with mutation in the genome stability regulators such as MRC1, SGS1, RAD53, SRS2, MEC1 and RTT101(11–14). The RRM3 gene has also been implicated in the suppression of the Ty1, Ty2 and Ty3 element (15, 16). Here in this paper we report the genetic interactions among the SAF1 and RRM3 genes. The simultaneous deletion of both gene leads to synthetic growth defects and genomic instability. The growth fitness of the double deletion strain was even further reduced when exposed to MMS and HU in comparison to single gene mutant or WT.

## Methods

### Strain and Plasmids

BY4741 (*Mata his3Δ1 leu2Δ0 met15Δ0 ura3Δ0*) *Saccharomyces cerevisiae* strain was used for the construction of the gene deletions. Strain JC2326 (*MAT-ura3, cir0, ura3–167, leu: his, his32*<u>*Ty1his3AI-270*</u>, *Ty1NEO-588*, *Ty1 (tub: lacs)-146)*was used for Ty1 retro-transposition assay (Table1).Following, plasmids, pFA6a-KanMX6, pFA6a-His3MX6, and pGADT7(Table 2) were used for PCR amplification of the deletion marker cassette containing 40 bp homology to the upstream and downstream of the target ORFs.

**Table 1:**
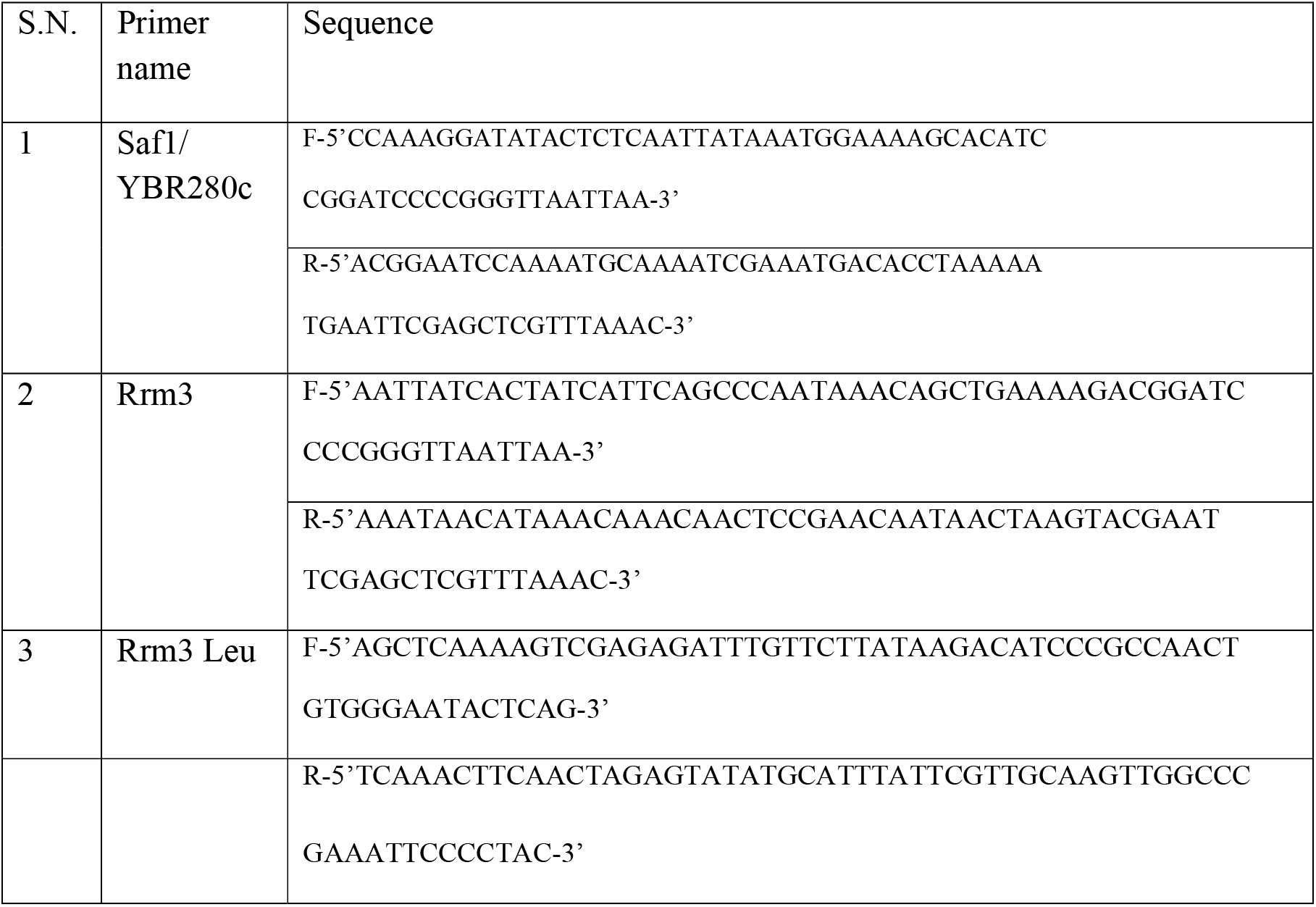
List of primers used for construction of deletion strains.

**Table 2:**
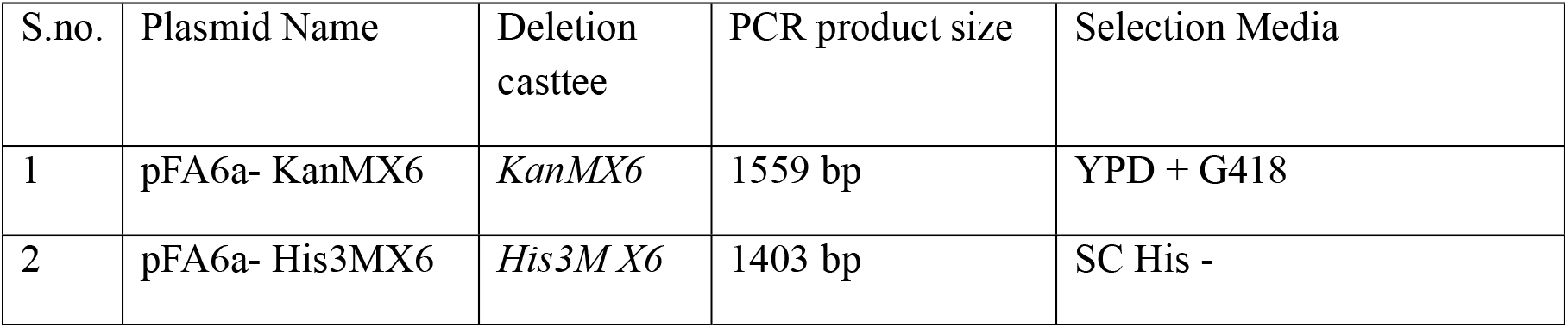
List of plasmids used for generating deletion cassette with selectable marker.

| S.no. | Plasmid Name | Deletion casttee | PCR product size | Selection Media |
| --- | --- | --- | --- | --- |
| 1 | pFA6a- KanMX6 | *KanMX6* | 1559 bp | YPD + G418 |
| 2 | pFA6a- His3MX6 | *His3M X6* | 1403 bp | SC His - |

**Table 3:**
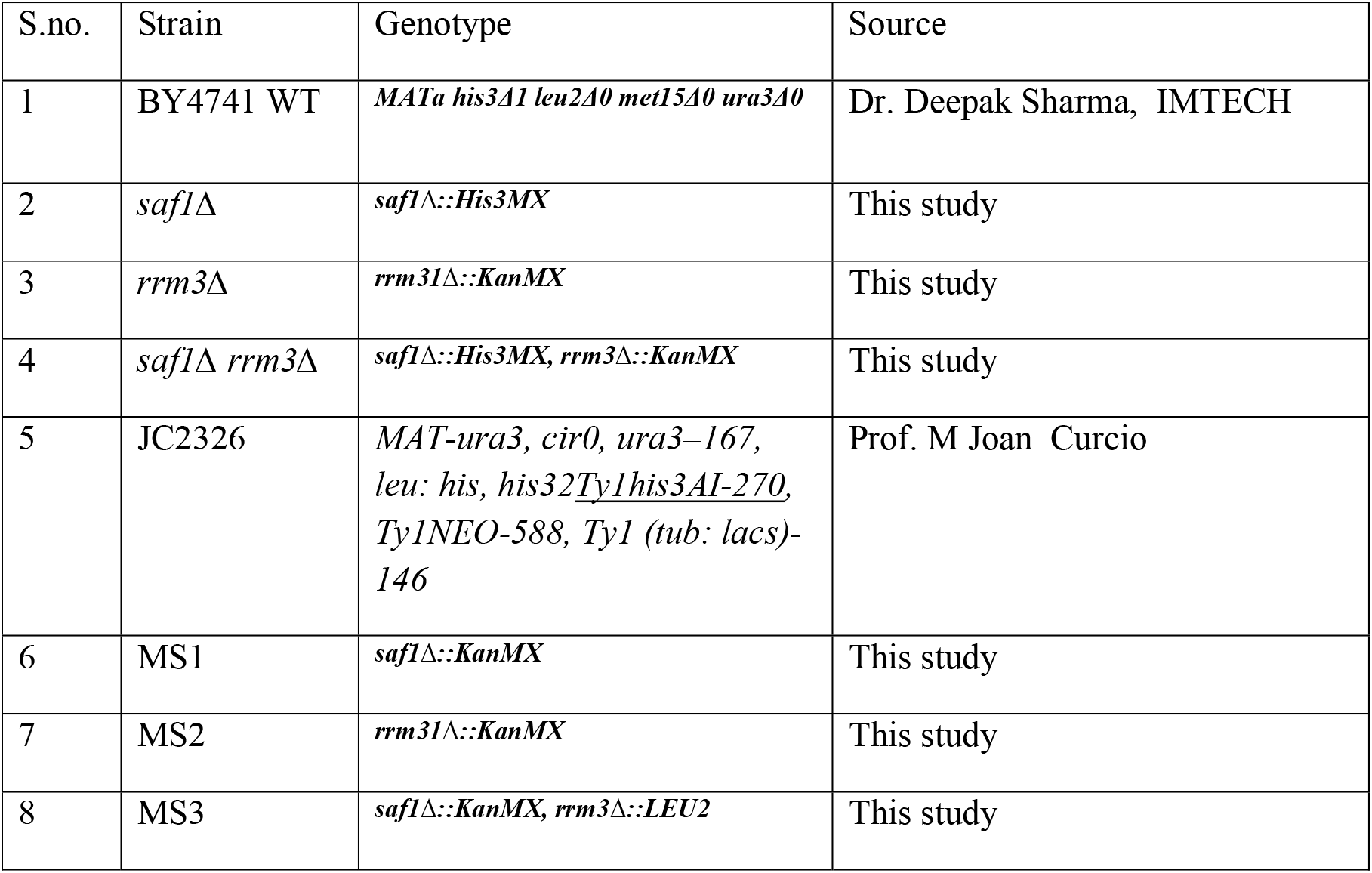
Yeast strains and their genotype used in the study.

### Growth Assay

Growth assessment of the WT and mutants was carried out as mentioned (17). Growth kinetics of strains was analyzed for 14 hrs by measuring optical density at 600nm using TOSHVIN UV-800 SHIMADZU spectrophotometer. The cultures were also streaked on the YPD plates followed by incubation at 30°C for 2-3 days.

### Phase Contrast Microscopy

For microscopy analysis, each strain was grown up to log phase in YPD medium at 30°C and imaged under Leica DM3000 microscope at 100X magnification.

### Semi Quantitative Growth Assay

To compare the growth fitness among the WT and mutants, spotting assay was performed. Wild type (BY4741) and its deletion derivatives strains (*saf1Δ,rrm3Δ* and *saf1Δrrm3Δ*) were grown in the 25 ml YPD (Yeast Extract 1% w/v, Peptone 2% w/v, dextrose 2% w/v) medium overnight at 30°C. The next day the cultures were diluted and grown in fresh YPD medium for 3-4hrs so as to reach early log phase (OD600 0.8-1.0). The cultures were equalized by OD at 600nm. A tenfold serial dilution was done and spotted (3µl) onto agar plates containing (YPD and YPD + HU and MMS). The plates were incubated at 30°C for 2-3 days and imaged.

### Nuclear Migration Assay using Fluorescence microscopy

For detection of nuclear migration defects, assay described in (18, 19) was adopted. The number of nuclei per cell in yeast strains was determined with nuclear binding dye (DAPI). Briefly, strains were grown to early log phase (OD_600_ ~ 0.8) at 30ºC. Yeast cells were washed with distilled water and suspended in 1X PBS (Phosphate Buffer Saline). Further, fixation was done by addition of 70% ethanol before DAPI staining. Cells were washed with 1X PBS then again centrifuged for 1 minute at 2500 rpm. DAPI stain (1mg/ml stock) to final concentration of 2.5µg/ml was added and incubated for 5 minutes at room temperature and visualized under UV light of fluorescent microscope with 100X magnification. A total of 200 cells were counted and grouped according 0, 1, 2 and multi nuclei per cell, more than two nuclei indicated the nuclear migration defect.

### Assay for Ty1 retro-transposition

To measure the Ty1 retro-transposition frequency the method mentioned in (16, 20) was followed. Briefly, single colony of JC2326 (reporter strain) and its deletion derivative (*saf1Δ* and *rrm3Δ* and *saf1Δrrm3Δ*) strains were inoculated into 10 ml YPD broth and was grown overnight at 30°C. The overnight grown cultures were again inoculated in 5ml YPD at 1:1000 dilutions and grown up to saturation point (144hrs) at 20°C. Then saturated culture was serially diluted and plated on minimal media (SD/His^−^ plates) followed by incubation at 30°C for 3-7 days. The frequency of appearance of His3^+^ colonies considered as measure of Ty1 retro-mobility.

### Statistical methods

Statistical significance of observations was determined using paired student t-test. P-value less than 0.05 indicated significant.

## Results

### The double mutant *saf1∆rrm3∆*showed synthetic growth defects

The null mutant of the SAF1 and RRM3 reported to be viable (21). Our data also showed the both single gene mutant (*saf1∆, rrm3∆*) as viable. The growth of the single gene mutant slightly reduced in comparison to WT (Figure 1A, C). The phase contrast imaging of the single gene mutant appeared normal just like WT however the *rrm3∆* mutant showed the enlargement of the cell (Figure 1 B).The growth characteristics of the double mutant *saf1∆rrm3∆*showed reduced growth phenotype in comparison to *saf1∆*, *rrm3∆* and WT when grown in YPD medium (Figure 1 (A, C). It has been reported previously that the growth of the *rrm3∆* is slower than the WT. The double mutant showed the synthetic growth defects as indicated by streaking. The growth kinetics of the *saf1∆rrm3∆* indicated that loss of both the gene leads to synthetic growth defects.

**Figure 1.**
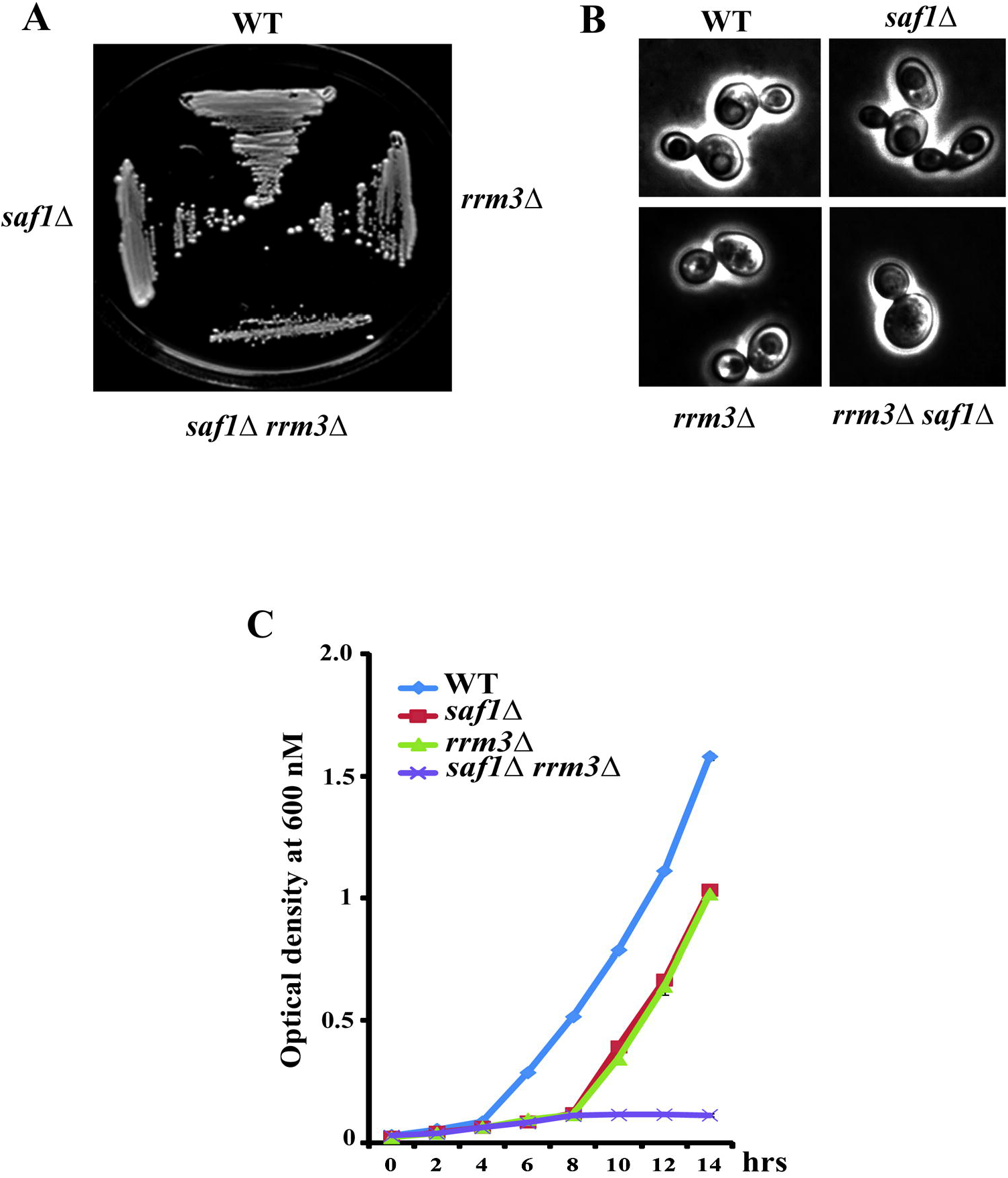
Comparative analysis of growth and morphology of WT, *saf1∆*, *rrm3∆*, *saf1∆rrm3∆* strains. A. Growth of streaked strains on YPD plates, streaked culture were incubated for 2-3 days at 30°C and then photographed. B. Phase contrast images of log phase cultures at 100X magnification using Leica DM3000. C. Growth kinetics of strains (WT, *saf1∆*, *rrm3∆*, *saf1∆rrm3∆*). Cells were collected every 2 hour period and cellular growth was measured by optical density (OD) at 600 nm using TOSHVIN UV-1800 SHIMADZU. The data shown represent the average of three independent experiments the error bars seen represent the standard deviation for each set of data.

### The *saf1∆rrm3∆* cells exhibits Nuclear Migration defects

The analysis of the nucleus using DAPI stain indicated increased percentage of cells showing multi-nuclei phenotype in *saf1∆rrm3∆* cellsin comparison to *saf1∆*, *rrm3∆*, and WT (Figure 2A). The multi-nuclei phenotype indicated the defect in the nuclear migration..

**Figure 2.**
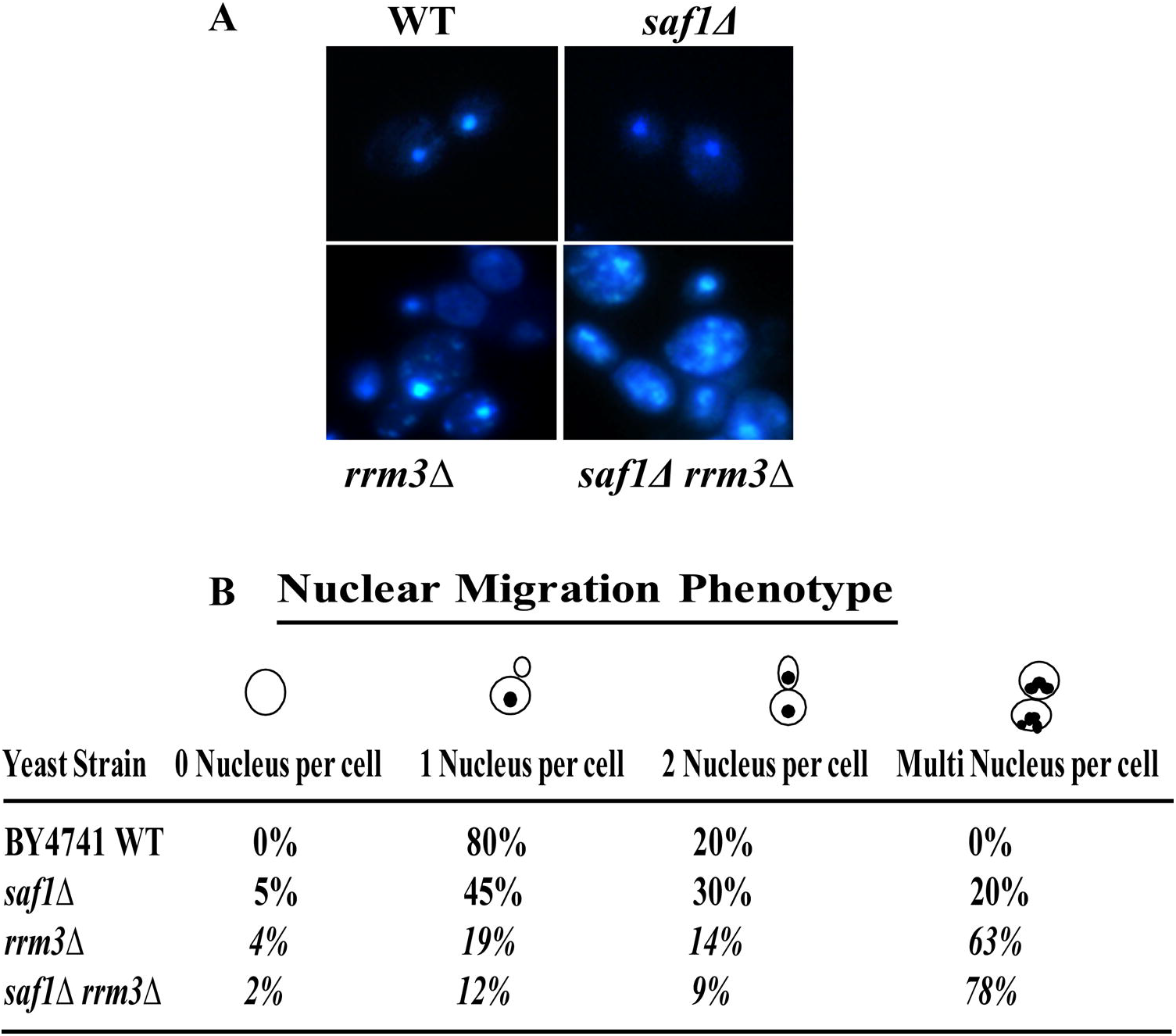
Assessment of nuclear migration phenotype using DAPI (4′,6-diamidino-2-phenylindole) staining of WT, *saf1∆*, *rrm3∆*, *saf1∆rrm3∆*. A. Representative Images of WT, *saf1∆*, *rrm3∆*, *saf1∆rrm3∆* cells showing the status of Nuclear DNA migration. The images were acquired at the 100X magnification using Leica DM3000 fluorescent microscope. B Table showing the percentage from the count of 200 cells as, 0, 1, 2 and multi nucleus in each strain, more than two nuclei indicate the nuclear migration defect.

### The *saf1∆rrm3∆*cells showed increased Ty1 retro–transposition

Ty1 elements are class of retro-transposons in the *S. cerevisiae* genome which remain in the dormant state due the host encoded genetic factors. In the absence of RRM3, the frequency of Ty1retro-transposition goes up as reported previously (15, 16). In this study the single gene mutant *saf1∆*showed nearly 8 fold whereas the *rrm3∆* showed the 85fold change in His+ prototroph formation in comparison to WT. However the absence of both the genes *(saf1∆rrm3∆)* leads to increased frequency (~ 180-fold) of Ty1-retro transposition in the comparison to WT cells. (Figure 3 A, B).

**Figure 3.**
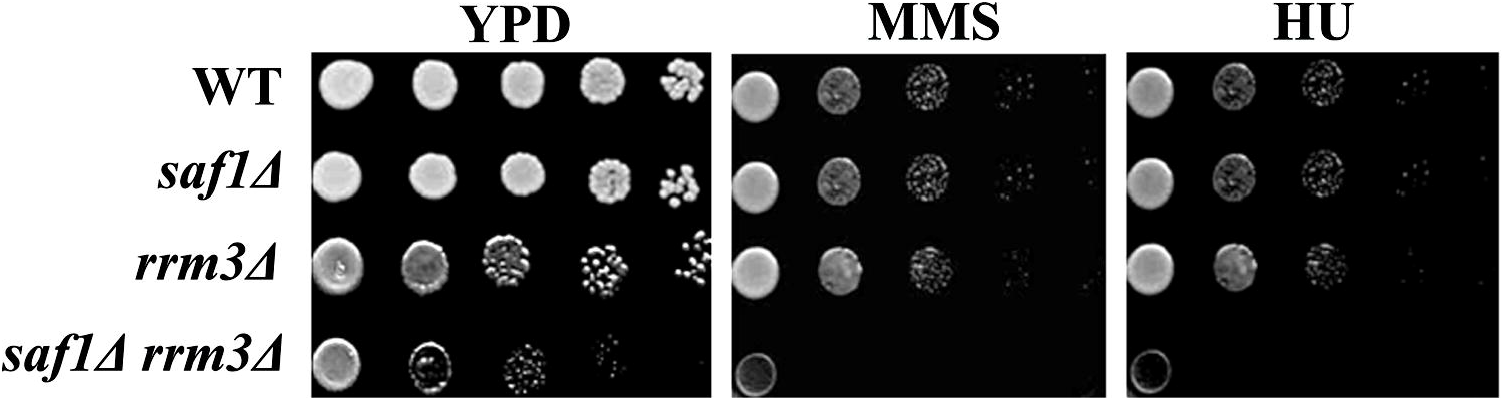
Assay for genome instability by measuring of *his3AI* marked Ty1 transposition frequency in WT, *saf1∆*, *rrm3∆*, *saf1∆rrm3∆*. A Images of plates showing the Ty1 transposition induced colonies on Synthetic Dropout (SD) plate lacking histidine. B. Bar diagram showing the frequency of Ty1his3AI transposition in each strain. The data shown represent the average of three independent experiments. The significance of transposition was determined by using two tailed t-test. P-value (p) less than 0.05 indicates significant difference and the symbol * represent to p<0.05..

### Loss of SAF1 and RRM3 leads to HU and MMS sensitivity

The hydroxyurea (HU) and methyl methanesulfonate (MMS) act as replication checkpoint inhibitor and DNA damaging agent respectively. The Rrm3 mutant reported to have slightly sensitive phenotype in the 200mM hydroxyurea (22). With regard to SAF1 mutant the data is not available. The semi-quantitative growth assay by spot analysis on YPD medium and YPD+genotoxic agents (MMS and HU) of the WT, *saf1∆*, *rrm3∆* and *saf1∆rrm3∆* indicated the extremely slow growth of the *saf1∆rrm3∆* when compared with the WT in presence of HU and MMS (Figure 4 A, B). The *saf1∆*, *rrm3∆* also showed the reduced growth in presence of the HU and MMS when compared with the WT.

**Figure 4.**
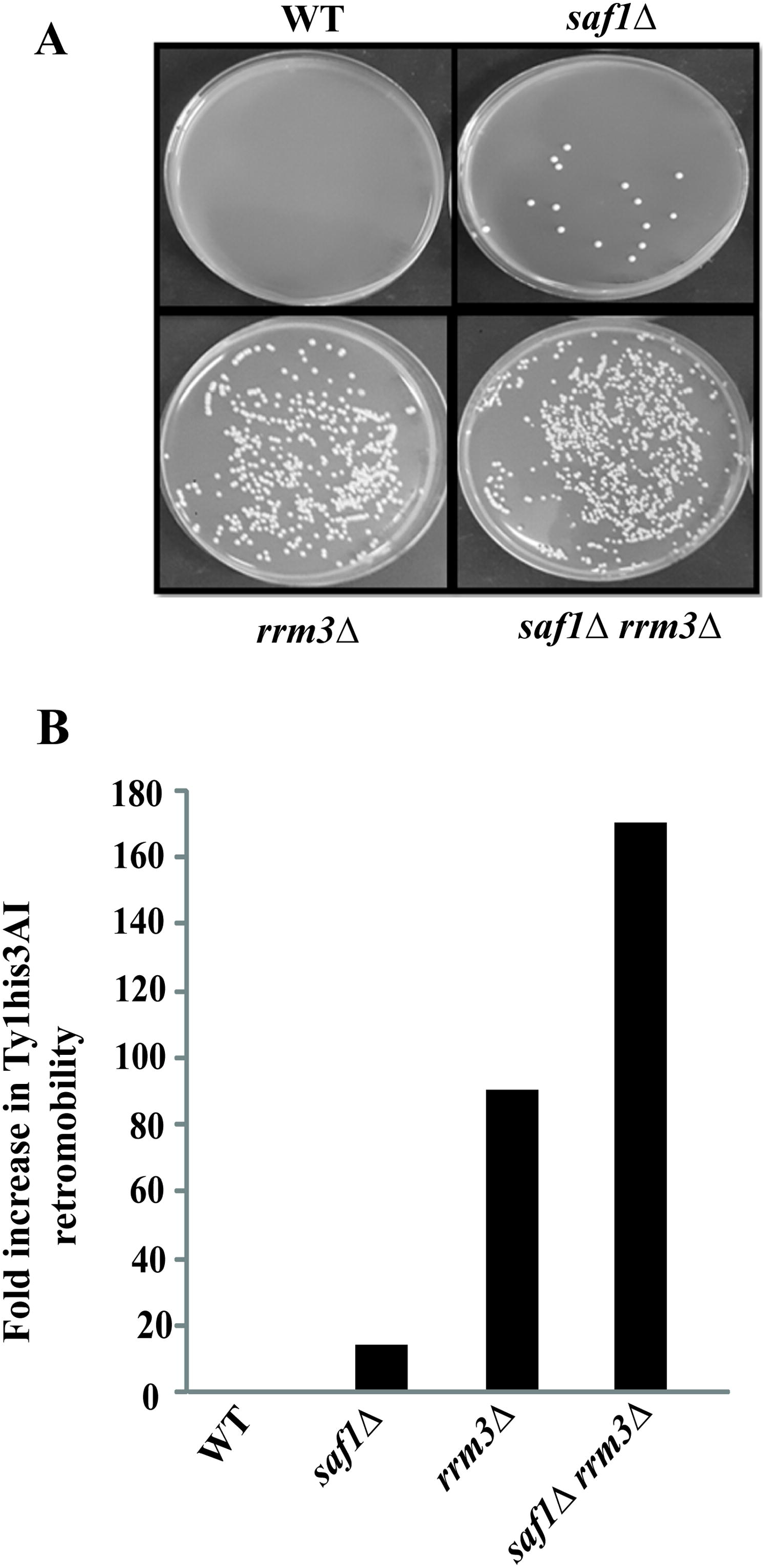
Comparative assessment of growth phenotype of WT, *saf1∆*, *rrm3∆ and saf1∆rrm3∆* strains in presence of 0.035% MMS and 200mM HU genotoxic stress causing agents. Each strain was grown to log phase and was equalized by O.D at 600 nm. The tenfold serially diluted samples of each strains spotted on YPD, YPD+stress agent containing agar plates. The result indicates that *saf1∆rrm3∆* double mutant exhibits the sensitivity in the presence of HU and MMS.

## Discussion

The multi-subunit enzymes complex, Skp1-Cul1-F-box (SCF) E-3, recruits the substrate or regulatory proteins through F-box motif containing protein for poly-ubiquitination and subsequent degradation by the 26S proteasome (23, 24). The Saf1 of *S.cerevisiae* is less characterized F-box protein which contains RCC1repeats (25). The Saf1 has been shown to involved in the phase transition through Aah1pdegradation during nutrient deprivation (25).In this paper we have investigated the genetic interaction among the SAF1and genome stability regulator RRM3. The search for the genetic interactions and their impact on phenotype is crucial for the overall understanding of the biological processes and functioning of molecular player. The investigations of the binary interactions are crucial in unearthing of the pathways for better understanding of the system biology. The null mutant of the SAF1 have been reported as viable (21)), exhibited the decrease in cell death (26) phenotype and resistance to histone deacetylase inhibitor CG-1521 drug (27). In our study also SAF1 null mutant showed the decreased growth in comparison to WT. The null mutant of RRM3 reported as viable (21) and showed the abnormal bud morphology (28). Our data with the *rrm3∆*, showed the phenotype as reported earlier. The RRM3mutant showed the slightly sensitive phenotype when grown in presence of 200mM HU (22) and MMS (0.01%) (29). In our study we observed that *rrm3∆* showed slightly reduced growth when grown in presence of the 200mM HU and 0.035% MMS. The retro-transposition frequency in *S.cerevisiae* genome reported to be elevated (16, 30) including Ty1 (31) and Ty3 elements (32) in the null mutant of RRM3. The genetic interaction studies carried out previously reported that SAF1 interact negatively with CDC 10, CDC11,CDC12, HYP2(33) and DDC1(34). The null mutant of SAF1 showed the synthetic growth defects with HSP82 (35), POL2 (36) and RTT109 (37). The rrm3∆ showed the synthetic growth defect with multiple genes such as CCR4, CCS1, CTF18, CTF4, CTF8, DCC1, MMS1,MRE11, PAP2, POP2, RAD51, RAD55, RAD57, RTT107, SRS2, SGS1(38)CTF8, DCC1, RAD52, RAD53 (39) in high throughput studies. However this is the first study which reports the genetic interaction among the SAF1 and RRM3 gene. Our data suggest that both the genes interact genetically to regulate the fitness of the cell and genome stability. The investigated genetic interaction may have implication for the cancer studies as there is higher degree of functional homology of Saf1p, Rrm3 with the mammalian HERC2 andPif1 helicase respectively. In conclusion, our investigation on binary genetic interaction with the SCF-3 ligase component F-box motif encoding gene SAF1 and genome stability regulator RRM3 suggest that loss of both the genes in *S. cerevisiae* reduces the growth fitness of the WT cells.

## Acknowledgment

We thank to Prof. M Joan Curio, Dr. Deepak Sharma, IMTECH, Dr. Ravi Manjithya, NCBS, India for strains and plasmids.

## Funding information

This work was supported by a grant (BT/RLF/Re-entry/40/2012) from the Department of Biotechnology, GOI, New Delhi to N.K.B who is recipient of the Ramalingaswami fellowship from DBT, New Delhi.

## Conflict of Interest

The authors declare that they have no conflicts of interest with the content of this article.

## Author’s contributions

NKB conceived and directed the study and wrote the paper with MS and VV.MS performed the experiments and analysed with NKB.VV provided the bioinformatics facility and analysed the data. All the authors reviewed the results and approved the final version of manuscript.

